# When blood is not thicker than water: Social information landscapes offset the food-fear trade-off in habitat selection decisions

**DOI:** 10.64898/2026.10.06.757038

**Authors:** Levi Newediuk, Zoe Melvin, Liam Thomas, Eric Vander Wal

## Abstract

1. Group foraging reduces the risks associated with foraging: larger groups dilute individual predation risk, allow group members to share the costs of vigilance, and provide a social structure for exchanging information about predators. The benefits of shared social information over simple safety in numbers should depend on the relationships among group members. Consequently, group composition should affect the selection of risky and safe habitats in social species.
2. We asked whether relationships among elk (*Cervus canadensis*) group members moving together influence their selection of risky habitats more than grouping alone. We tested three hypotheses. The *familiarity hypothesis* predicts that elk with close social ties are most likely to select risky habitats together. The *inclusive fitness hypothesis* predicts that the closest genetic relatives are most likely to select risky habitats together. The *dilution hypothesis* predicts that familiarity and genetic relationships do not influence habitat selection, with all group foraging benefits derived from proximity to others.
3. We used integrated Step Selection Analysis to test whether the identity of the closest group members influenced habitat selection. We tested whether the relationship of the nearest elk—either genetic or social—incentivized selection of risky habitat more than proximity to the nearest elk.
4. Group foraging clearly reduced risk, with elk more likely to select risky habitat when near others. Elk were also more likely to select risky habitats when near closer relatives, albeit with considerable variation among individuals. However, familiarity had a large and consistent effect: elk were far more likely to select risky habitats when near socially familiar group members.
5. Our results suggest that information shared within foraging groups, especially between socially familiar individuals, may lower the risks associated with habitat selection. Similarly, the benefits of reliable social information for reducing risk likely incentivizes the formation of stronger social ties within foraging groups. This positive feedback between social familiarity and habitat selection could thereby increase access to profitable habitats that are otherwise too risky, cementing what we term the *social information landscape* as a key driver of movement and habitat selection.

## Introduction

Foraging is an essential activity that is also inherently costly. Predators search for prey in the same places where prey search for forage (Sih, 2005), increasing the risk of predation while prey also lose foraging efficiency to vigilance (Cowlishaw et al., 2004; Lima & Dill, 1990). Habitat selection decisions, therefore, hinge on whether accessing the rewards of high-quality foraging habitat is sufficient to pay for the associated costs. From a movement ecology perspective, these decisions emerge from the interaction between an individual’s internal state and its external environment (Nathan et al., 2008). Energetic needs, for example, modify the value of searching for food relative to avoiding predators (Houston et al., 1993), while the distribution of food and predators determines where costs and rewards are encountered when individuals move (Laundré et al., 2001). However, social prey animals also navigate these individual decisions within social groups (Webber et al., 2023). Social foraging dilutes individual predation risk and spreads the responsibility for vigilance across the group (Pulliam, 1973), allowing individuals to forage in places they might otherwise avoid (Beauchamp, 2013). The extent to which social groups reduce risk may depend, however, on which group members are present and the benefits they provide to offset the costs of foraging.

Social foraging lowers the risk to individual foragers in two ways. First, foraging in groups spreads risk over multiple group members, numerically diluting the risk of predation to any single individual (Lehtonen & Jaatinen, 2016; Lima & Dill, 1990). This risk reduction increases the net benefit of foraging in risky places. Second, social foraging allows group members to share information about predators through alarm signals and other cues that warn group members of danger (Dall et al., 2005; Favreau et al., 2010; Gil et al., 2017; Pays et al., 2013). Access to this collective information allows individuals to detect predators sooner and respond more effectively (McCormick et al., 2026; Nadler et al., 2021), further balancing the rewards and costs of foraging. While information-sharing and numerical dilution likely operate simultaneously to lower risks whenever individuals forage in groups, their mechanisms differ: risk-dilution depends only on the number and proximity of nearby individuals, whereas the benefits of social information depend on whether nearby group members indeed possess information that lowers the risk of predation.

Not all social partners are equally valuable sources of information. Individuals under risk exhibit more effective antipredator responses when they forage in tighter-knit social groups with preferred partners (Chivers et al., 1995; Hasenjager & Dugatkin, 2017; Heathcote et al., 2017), suggesting familiarity reinforces the reliability of social information about risk. Information from relatives may be particularly valuable because warning relatives can increase the signaller’s inclusive fitness (Sherman, 1977), providing an evolutionary incentive for reliable information exchange. Consistent with this possibility, individuals are often most responsive to information about predators from relatives (Atherton & McCormick, 2017; Colombelli-Négrel & Evans, 2017), and many social species prioritize persistent associations between relatives and familiar individuals (Kurvers et al., 2013; Wild et al., 2024). These patterns suggest that the benefits of social information can extend beyond immediate predation risk to shape the long-term social structures animals forage within, and the habitats they select as a result.

Consequently, habitat selection should depend not only on the number of group members nearby, but also on their relationship with the focal individual. Since risk varies spatially, understanding the relative importance of familiar individuals versus genetic relatives for shaping habitat selection decisions could help reveal how the nexus of spatial risks, social networks and group composition—hereafter, the *social information landscape*—contributes to much broader space-use patterns at the population and species level. Such a connection would also position the social information landscape as an essential component of the movement ecology framework (*sensu* Nathan et al., 2008).

We tested whether elk (*Cervus canadensis*) group members vary in their capacity to balance the rewards and costs for foraging individuals, lowering the threshold level of safety required to accept risk while foraging. In our study system, elk have access to open, high-risk but productive habitats and closed, safer habitats with lower productivity. We hypothesized that more elk nearby would reduce risk in open habitats through dilution, but if elk also acquire information from group members, the magnitude of the benefit should depend on the identity of the group members. Within this overarching hypothesis, we tested three separate hypotheses:

1. The *dilution hypothesis*: Any nearby individual should lower the safety threshold for habitat selection regardless of identity (Lima & Dill, 1990), increasing the likelihood that an elk closer to another individual would select a risky, open habitat rather than a closed habitat (Prediction 1).
2. The *familiarity hypothesis*: Nearby individuals who are also familiar should further lower the safety threshold by providing reliable social information, making it yet more likely that an elk nearby a familiar individual would select for open habitat (Prediction 2).
3. The *inclusive fitness hypothesis*: Kin have an additional incentive to provide reliable information about risk to their relatives. These individuals should lower the safety threshold to the greatest extent when nearby, making it most likely that elk near their relatives would select for open habitat (Prediction 3).

## Methods

### Biotelemetry data

Our study population consists of approximately 150 elk in southeast Manitoba, Canada (49.13, −96.56). The elk inhabit a largely agricultural landscape, approximately 40% of which is forested. The remaining landscape is open. While some wetland and natural prairie remain, the open habitat is largely cropland and pastureland, providing a productive source of forage for the elk that is also risky due to its exposure to natural predators (e.g., black bears, wolves) and humans. We captured 18 female elk, approximately 12% of the population, in February 2019 using a net gun fired from a helicopter. Each animal was fitted with an Iridium satellite collar (Vertex Plus 830 g, VECTRONIC Aerospace GmbH, Berlin, Germany) programmed to collect 30-minute GPS locations from 1st May to 31st August 2019 and 2020. All captures and sampling were conducted in accordance with approved animal care protocols (Memorial University of Newfoundland AUP #19-01-EV).

We used GPS location data collected from the collared elk to assess the relationship between social behaviour, habitat selection, genetic relatedness and movement. We screened the data and removed two-dimensional fixes, biologically implausible step lengths, and erroneous spikes between points (Bjørneraas et al., 2010). After screening, there was a mean of 48,233 (sd = 9,510) fixes per animal per year.

### Landscape classification process

We reclassified all habitat in our study system as either open or closed based on Agriculture and Agri-Food Canada (AAFC)’s 2019 and 2020 annual crop inventories. These spatial products provide 30-m resolution raster tiles of 60 crop types in addition to natural habitats derived from Landsat satellite imagery. We reclassified the AAFC’s shrubland, coniferous, and broad-leaf forests, or anything providing concealing cover for elk, as closed habitat. We classified all other available habitats as open, including all crops, urban and developed habitats, grasslands, wetlands, and pasture. We assumed open water was unavailable. Our earlier research based on activity timing and hormone production corroborates that open habitats are both riskier and more rewarding than closed habitats (Newediuk et al., 2026).

### Social network covariates

We used the methods described in Webber et al. (2024) to identify social associations based on the elk GPS location data using the *spatsoc* v0.2.13 (Robitaille et al., 2019) and *igraph* v2.3.3 (Csárdi & Nepusz, 2006) packages. Briefly, we considered individuals associated if their GPS locations were no more than 50 m from one another within 5 minutes. We used these associations in a moving window approach to generate weekly social networks (Webber et al., 2024) from May through August. The social network edges represent the frequency of associations between individuals, and nodes are individuals. We used the social network edges to calculate a modified simple ratio index (SRI) for all dyads, a single measure reflecting the number of repeated associations between individual elk (Webber et al., 2024).

### Genetic relatedness estimates

We estimated pairwise genetic relatedness between individual elk using microsatellite data. We collected a whole-blood sample from elk at the time of capture, which we sent to the Natural Resources DNA Profiling & Forensics Centre in Peterborough, Ontario, for DNA extraction and downstream genetic analysis. Extractions were performed using the DNeasy 96 Blood and Tissue kit extraction protocol (Qiagen, Germany). Individuals were then identified by amplifying and sizing nine microsatellite loci using GeneMarker software (SoftGenetics, USA).

We tested all individuals for pairwise genetic relatedness using the program *COANCESTRY* (Wang, 2011). Our measures of genetic relatedness are relative rather than absolute because we used the same samples for calculating relatedness as we used for the reference population (Wang, 2002). Several relatedness estimators are available in *COANCESTRY*; we used the Wang (2002) estimator because of its effectiveness on small datasets and its robustness to including related individuals in both the reference and sample populations (Wang, 2017). We ran bootstrapping of 100 for 1,000 interactions.

### Statistical analysis

We used integrated Step-Selection Analysis (iSSA), a habitat selection model, to test the influence of the identity and proximity of nearby elk on their selection of open habitat. In iSSA, the steps an animal takes—the straight-line distances between one GPS location and the next—are compared to “available” steps the animal could have taken from the same starting location (Avgar et al., 2016). The model estimates the density of used steps relative to available steps according to environmental covariates. It is often analyzed as an inhomogeneous point Poisson process to make it possible to incorporate random effects for multiple individuals (Muff et al., 2020). Additionally, covariates are estimated for the step length and turn angle between each subsequent step to account for the impact of the movement process on habitat availability (Avgar et al., 2016).

We used the *amt* package v0.3.1.0 (Signer et al., 2018) to generate a movement track for each elk, separating points into unique movement bouts when separated by more than 3 minutes. We also generated 10 available points for each used location, drawing random step lengths from a gamma distribution, and turn angles from a von Mises distribution parameterized using the characteristics from the elk movement data (Avgar et al., 2016).

We assigned social and habitat information to each used and available location point. We calculated the distance of the elk nearest the focal individual at the start of each step as a measure of nearest-neighbour distance. We further assigned the start of each step a genetic relatedness estimate and SRI based on the identity of the nearest neighbour, to assess the influence of group member genetic relatedness, familiarity, and proximity on the selection of open habitat relative to closed habitat. We then used the *raster* package v3.6-32 (Hijmans, 2025) to extract the reclassified landscape data, assigning the used and available locations a habitat type, either open or closed.

We fit three separate models to test our hypothesis about social information and the identity of group members. Each model included as covariates habitat type, either open or closed, at the end of each step and one social covariate, either the natural log of the nearest-neighbour distance to the closest elk at the start of its step or the natural log of the SRI or pairwise genetic relatedness index with respect to the nearest elk at the start of the step. This resulted in three models, one for each of our three predictions. We also included an interaction between the social covariate and habitat type to test the influence of nearby group member identity on the focal elk’s selection for open habitat relative to closed habitat. Additionally, we included a random intercept for elk ID in each model to account for repeated observations from individuals and a random slope for the interaction between the social and habitat type covariates. Finally, we included covariates for the natural log of the step length and the cosine of the turn angle to account for the movement process, and a random intercept for movement bout ID.

In addition to our models for hypothesis testing, we fit a baseline model without any social covariates or interaction, hereafter the ecological null model. A full list of models and covariates is in Table S1. We fit all models using the *glmmTMB* package v1.1.14 (Brooks et al., 2017) using a Poisson error distribution with a log link function and large fixed variances for the random effects (Muff et al., 2020). We set the random effect variances at 10,000.

We used likelihood ratio tests to compare the influences of proximity to, familiarity with, and pairwise genetic relatedness to the nearest elk at the start of each step on habitat selection at the end of the step. We tested the fit of these models against the ecological null model. We then used relative selection strength (RSS; Avgar et al., 2017) to visualize the strength of selection for a point in open habitat relative to closed habitat along a gradient of distance to the nearest neighbour, ranging from 0 m to 2 km; genetic relatedness to the nearest neighbour, ranging from an index value of -1.0 to 0.5; and SRI, ranging from 0 to 1. RSS is an approach to interpret the effects of covariates in habitat selection models across a gradient, while holding constant all other effects in the model (Avgar et al., 2017). We conducted all data cleaning and analysis in R v4.5.3 (R Core Team, 2026).

## Results

### Social covariates

Our social covariates and pairwise genetic relatedness estimates indicated that the distribution of pairwise genetic relatedness to elk at the start of steps was roughly uniform to slightly left-skewed (Supplementary Figure 1), while nearest-neighbour distances and SRI were right-skewed (Supplementary Figures 2 & 3). Genetic relatedness indices ranged from -1.04 to 0.83 with a median of 0.09, nearest-neighbour distances ranged from 0 to 24,267 m with a median of 740 m, and SRI ranged from 0 to 1 with a median of 0.01.

### Model results and comparisons

The ecological null model showed that elk displayed a slight avoidance of open habitat relative to closed habitat (Table 1). All three social models indicated that the nearest elk and its proximity at the start of a step altered the selection for open over closed habitat at the end of the step (Table 1). For example, an elk starting its step 5 m from another elk selected open habitat over closed habitat approximately 22% more strongly than an elk starting 500 m from another elk (*dilution effect model*; Figure 1). When starting near a closer genetic relative (75^th^ percentile genetic relatedness) versus a more distant relative (25^th^ percentile genetic relatedness), elk selected open over closed habitat approximately 9% more strongly (*inclusive fitness model*; Figure 2). When starting near a highly familiar individual (75**^th^** percentile SRI) versus an unfamiliar individual (25^th^ percentile SRI), elk selected open habitat over closed habitat approximately 70% more strongly (*familiarity model*; Figure 3).

**Figure 1.**
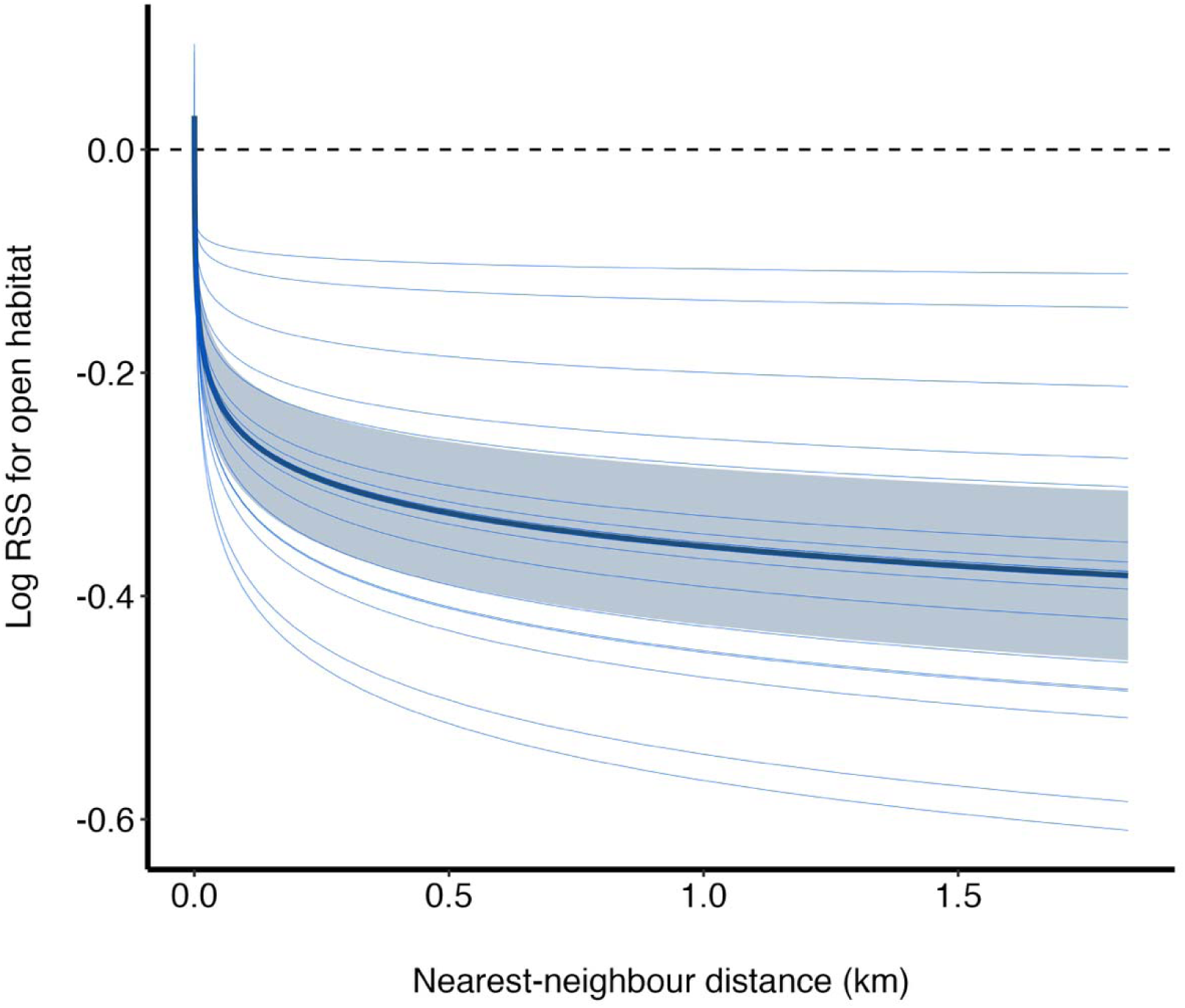
Log relative selection strength (RSS) for a point in open habitat relative to closed habitat across a gradient of distance to the nearest elk. The dotted line represents the point at which the preference for closed over open habitat begins to reverse, and there is a proportionate increase in selection for open habitat. The light blue lines show individual elk responses (n = 18), while the dark blue line shows the population-level response, including the 95% confidence interval of the RSS estimate (light blue envelope). The graph shows that relative selection for open habitat relative to closed habitat drops sharply when there are more than a few metres between individuals.

**Figure 2.**
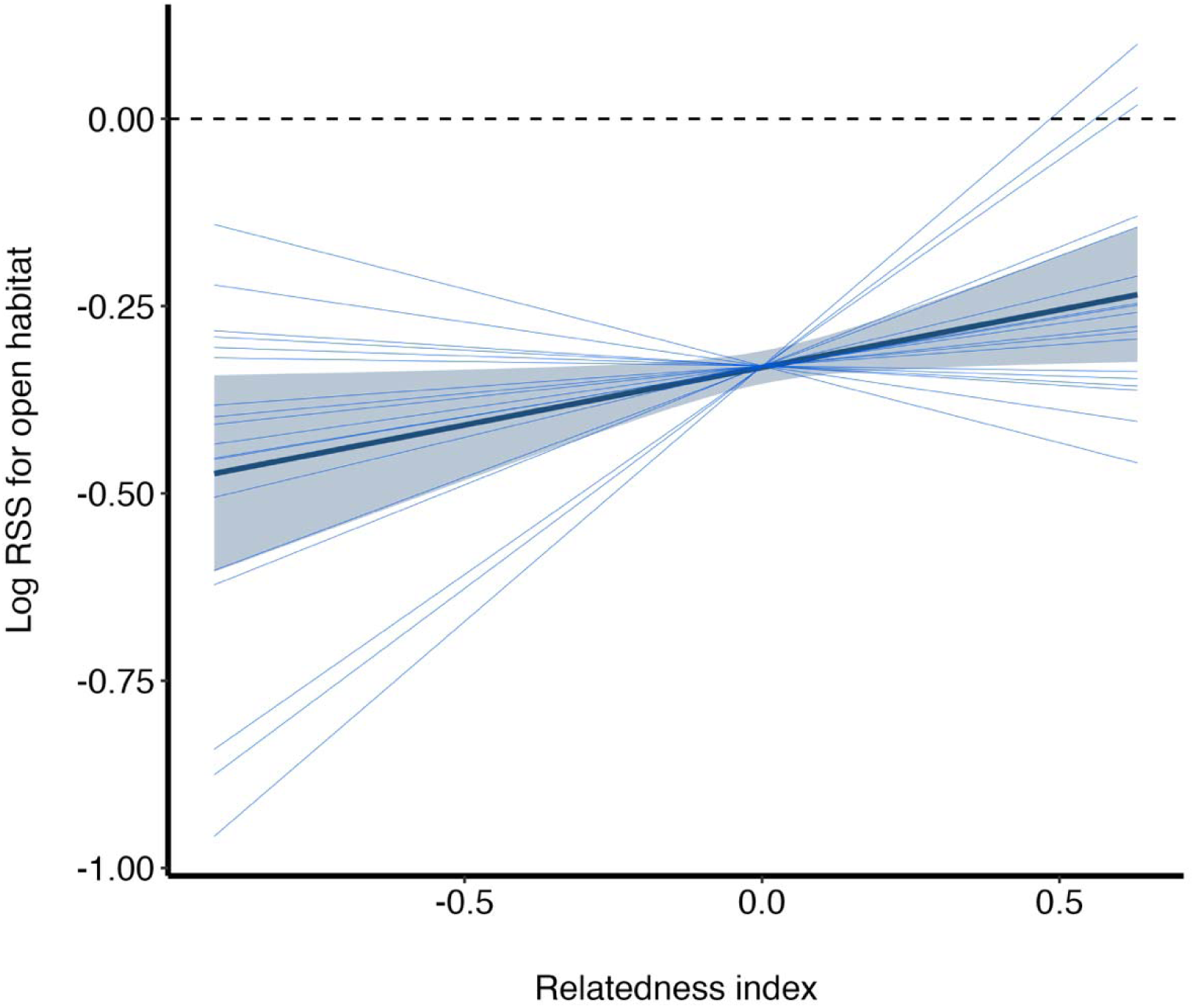
Log relative selection strength for open habitat relative to closed habitat across a gradient of genetic relatedness to the nearest elk. The dotted line represents the point at which the preference for closed over open habitat begins to reverse, and there is a proportionate increase in selection for open habitat. The light blue lines show individual elk responses (n = 18), while the dark blue line shows the population-level response, including the 95% confidence interval of the RSS estimate (light blue envelope). The graph shows a slight increase open habitat relative to closed habitat when the nearest elk is a closer relative for some individuals, with others experiencing a modest effect or the opposite response.

**Figure 3.**
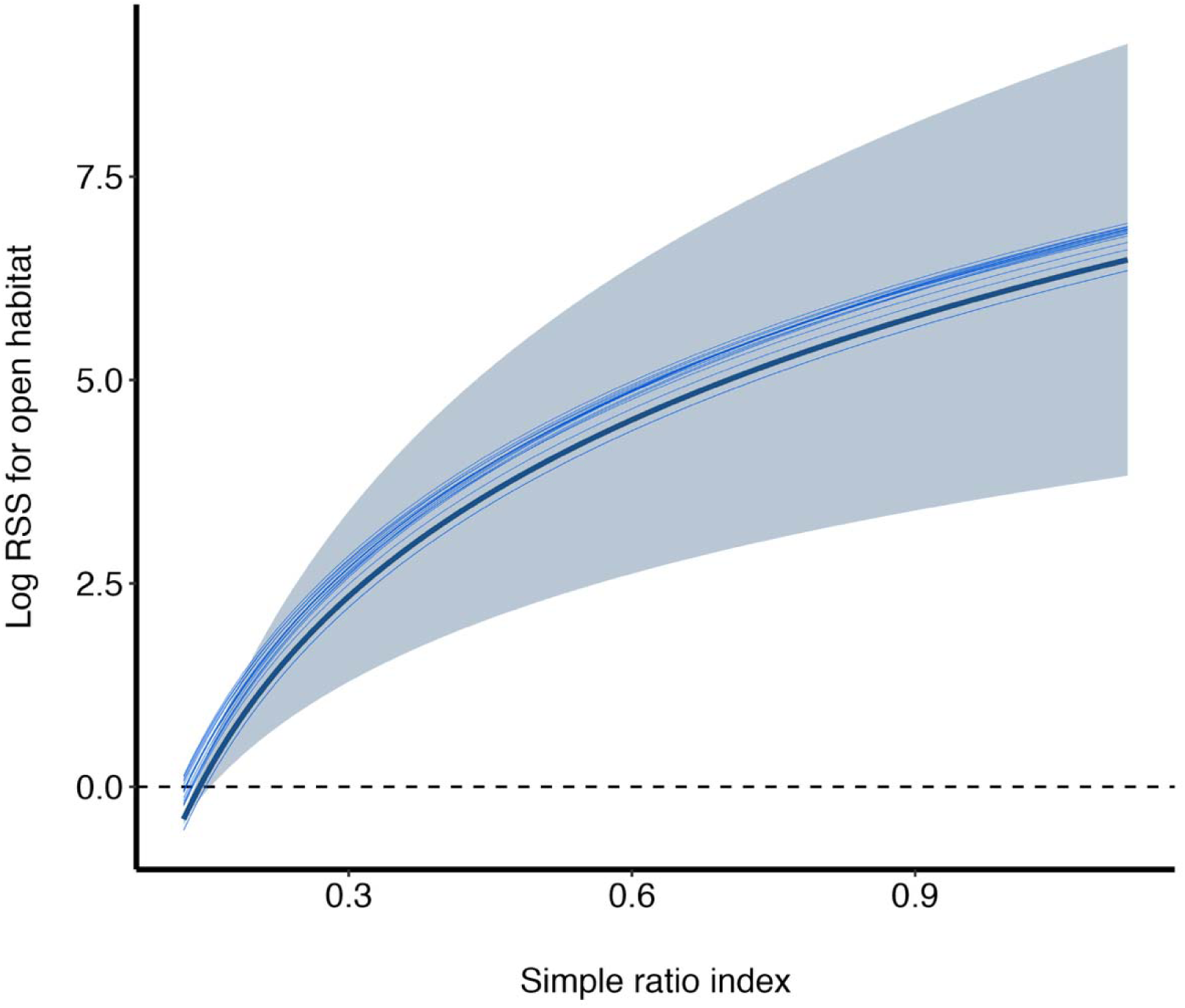
Log relative selection strength for open habitat relative to closed habitat across a gradient of simple ratio index (SRI) with respect to the nearest elk. The dotted line represents the point at which the preference for closed over open habitat begins to reverse, and there is a proportionate increase in selection for open habitat. The light blue lines show individual elk responses (n = 18), while the dark blue line shows the population-level response including the 95% confidence interval of the RSS estimate (light blue envelope). The graph shows that selection for open habitat relative to closed habitat increases sharply when the nearest elk has a closer social association.

**Table 1.** Coefficient estimates, standard errors (SE), and 95% confidence intervals (CI) for integrated step selection analyses estimating selection for open habitat relative to closed habitat. All models also include as covariates log(step length), cos(turn angle), and random intercepts for individual elk and individual movement bouts (separated by ≥ 3 minutes). Additionally, the models with social covariates (simple ratio index, SRI; nearest-neighbour distance, NN distance; genetic relatedness) include random slopes for the interaction between habitat type and the social covariate. Sample sizes for all models are n = 18 individuals and 1,433,288 step observations in 130,931 bouts. A full list of model covariates is in Supplementary Table S1.

| Covariate | $\beta$ estimate (± SE) | 95% CI |
| --- | --- | --- |
| **Familiarity model** |  |  |
| Habitat (open/closed) | $6.11 \pm 1.28$ | 3.60, 8.62 |
| Habitat x $\log(\text{SRI})$ | $3.13 \pm 0.62$ | 1.91, 4.35 |
| **Dilution model** |  |  |
| Habitat (open/closed) | $-0.06 \pm 0.02$ | -0.11, -0.01 |
| Habitat x $\log(\text{NN distance})$ | $-0.04 \pm 0.01$ | -0.05, -0.03 |
| **Inclusive fitness model** |  |  |
| Habitat (open/closed) | $-0.33 \pm 0.01$ | -0.35, -0.31 |
| Habitat x relatedness | $1.54 \pm 0.07$ | 0.01, 0.29 |
| **Ecological null model** |  |  |
| Habitat (open/closed) | $-0.31 \pm 0.001$ | -0.33, -0.29 |

Likelihood ratio tests indicated that familiarity, dilution, and relatedness models were all significant improvements over the ecological null model. When considering the identity of the nearby individuals, the familiarity model was most likely compared to the ecological null model ( ^2^(3) = 371,508, p < 0.0001). The dilution effect model was next most likely ( ^2^(3) = 323.8, p < 0.0001), followed by the inclusive fitness model ( ^2^(3) = 104.2, p < 0.0001).

## Discussion

Foraging entails rewards and costs, and animals have ecological strategies to ensure a net reward. Group foraging dilutes the individual risk of predation and frees up time for foraging by spreading the responsibility for vigilance across group members (*dilution hypothesis*; Beauchamp, 2013; Pulliam, 1973). Group structures also facilitate shared information about risk (Gray & Webster, 2023); we hypothesized that the reliability of information should depend on the identity of the individual conveying it. We tested whether the identity and proximity of group members determined how much their proximity balanced the costs of foraging. As expected, we found that female elk were more likely to select risky, open habitat the closer they were to any other elk before moving, indicating a risk-dilution effect of group foraging. Group members that are relatives should moderate risk most, as they have an incentive to provide reliable information to their relatives to increase their inclusive fitness (*inclusive fitness hypothesis*). As predicted, we found that selection for open habitat increased when genetic relatives were nearby. However, nearby familiar elk increased selection for open habitat seven times more than relatives and to a greater extent than proximity alone, suggesting that social ties within foraging groups are sufficient to lower risk (*familiarity hypothesis*). By facilitating access to information about risk, familiar social partners fundamentally changed the perceived costs associated with productive habitats, and with that, the habitat selection decisions of individual elk. Our results suggest that habitat selection for social animals is not only simple function of group size and configuration but also emerges as a product of the surrounding social information landscape.

What about familiarity might incentivize risky habitat selection more than kinship, despite the inclusive fitness benefits of sharing reliable information with relatives? We suggest that the importance of social information likely depends not only on the reliability of the signaller but on the degree to which the information being conveyed is relevant to the recipient. Social ungulates often segregate into sex and age classes because their energy and safety requirements align (Bon & Campan, 1996; Ruckstuhl & Neuhaus, 2002). Female elk, for example, form close-knit social groups with other females during summer because they experience similar predation pressure and benefit from shared vigilance and calf defence (Geist, 1982), with growing familiarity as they interact frequently. Aligned interests may be a strong incentive to maintain close associations when facing risk, possibly to a greater extent than many close relatives from different age and sex classes. Consequently, information from familiar individuals likely mitigates risk more consistently than information shared among relatives.

Familiar individuals might also mitigate risk more consistently because they have access to preferred social information not exclusively available to kin. In social networks, central individuals act as hubs for interaction (Wey et al., 2008). For social groups that engage in risky foraging, some individuals may become central because they have exclusive access to reliable information about risk. Studies of shoaling fish, for example, have shown that startle responses are consistently initiated by the same central individuals in the group (Marras & Domenici, 2013). Followers then gain a fitness benefit by attending to information from these group leaders (Ioannou et al., 2019). Leader-follower interactions suggest that by virtue of their central positions, some individuals in social groups are also most familiar to all group members. If familiarity coincides with access to information, individuals likely perceive less risk and consequently alter their habitat selection when near any central familiar individuals, regardless of genetic relatedness.

Although we found familiarity was most influential for selecting risky habitat, this is not necessarily evidence that genetic relatedness is unimportant. Genetic relatedness and familiarity are often closely linked (Doran et al., 2019; Lee-Jenkins & Godin, 2013), making their effects difficult to separate. When relatives associate more often than unrelated individuals, associations attributed to relatives may reflect familiarity. Consequently, familiarity may already capture much of the information benefit of kinship, leaving little additional incentive to obtain information from kin beyond that which is common in repeated social interactions, and potentially less incentive when relatives have conflicting ecological motivations. The overlap between kinship and familiarity is likely particularly pronounced in our study population, where female elk form stable matrilineal social groups (Geist, 1982). Although our genetic relatedness estimates were relative, preventing us from characterizing the exact genetic relationships between individuals, they estimates were skewed toward the higher end of relatedness (Supplementary File 1; Supplementary Figure 1). The likely low resolution of genetic relatedness in our study population suggests we had limited ability to detect the role of kin selection for the reliability of social information. Disentangling familiarity from genetic relatedness through experimental manipulation may therefore be key to determining the extent to which information from kin provides any additional benefit over information from social partners.

## Conclusions

Our results highlight that the social information landscape is a key factor in social herbivores making habitat selection decisions, particularly in balancing the costs and benefits of foraging. Other dynamic landscapes tied to foraging, such as the landscape of fear (Laundré et al., 2001) and energy landscapes (Wilson et al., 2011), are already established and embedded in the movement ecology framework. However, both fear and energy acquisition depend greatly on access to social information, meaning the social information landscape is potentially foundational to many of the insights developed in movement ecology over the past few decades. Based on our results, in many instances access to social information likely also surpasses simple risk dilution in its influence over movement and habitat selection.

The importance of social information we demonstrate here supports the view that habitat selection models should move beyond simple measures of population density and group size towards covariates that reflect the dynamic socioecological environment (Webber & Vander Wal, 2018). A more recent and growing body of research integrating social and habitat influences has revealed new insights showing that social environments indeed influence habitat selection decisions (Atkinson et al., 2025; Hendrix et al., 2024; Webber et al., 2024). However, we have yet to parse the influence of individual group composition and the information contained therein on habitat selection. We suggest that insights into the drivers of habitat selection will continue to grow as the social landscape—and more specifically, the social information landscape—becomes a more formalized concept in movement ecology.

## Data availability

Data and code in progress are both available from https://github.com/ljnewediuk/elk_social_issa.git. We intend to publish the final version of the repository as a Zenodo release upon acceptance of the paper.

## Supporting information

Relatedness estimates

