## Supplementary Materials for "When blood is not thicker than water: Social information landscapes offset the food-fear trade-off in habitat selection decisions"

**Supplementary Figure 1:** Distribution of pairwise genetic relationship estimates at the start of movement steps

**Supplementary Figure 2:** Distribution of nearest-neighbour distances at the start of movement steps

**Supplementary Figure 3:** Distribution of simple ratio indices at the start of movement steps

**Supplementary Table 1:** Full list of covariates in models

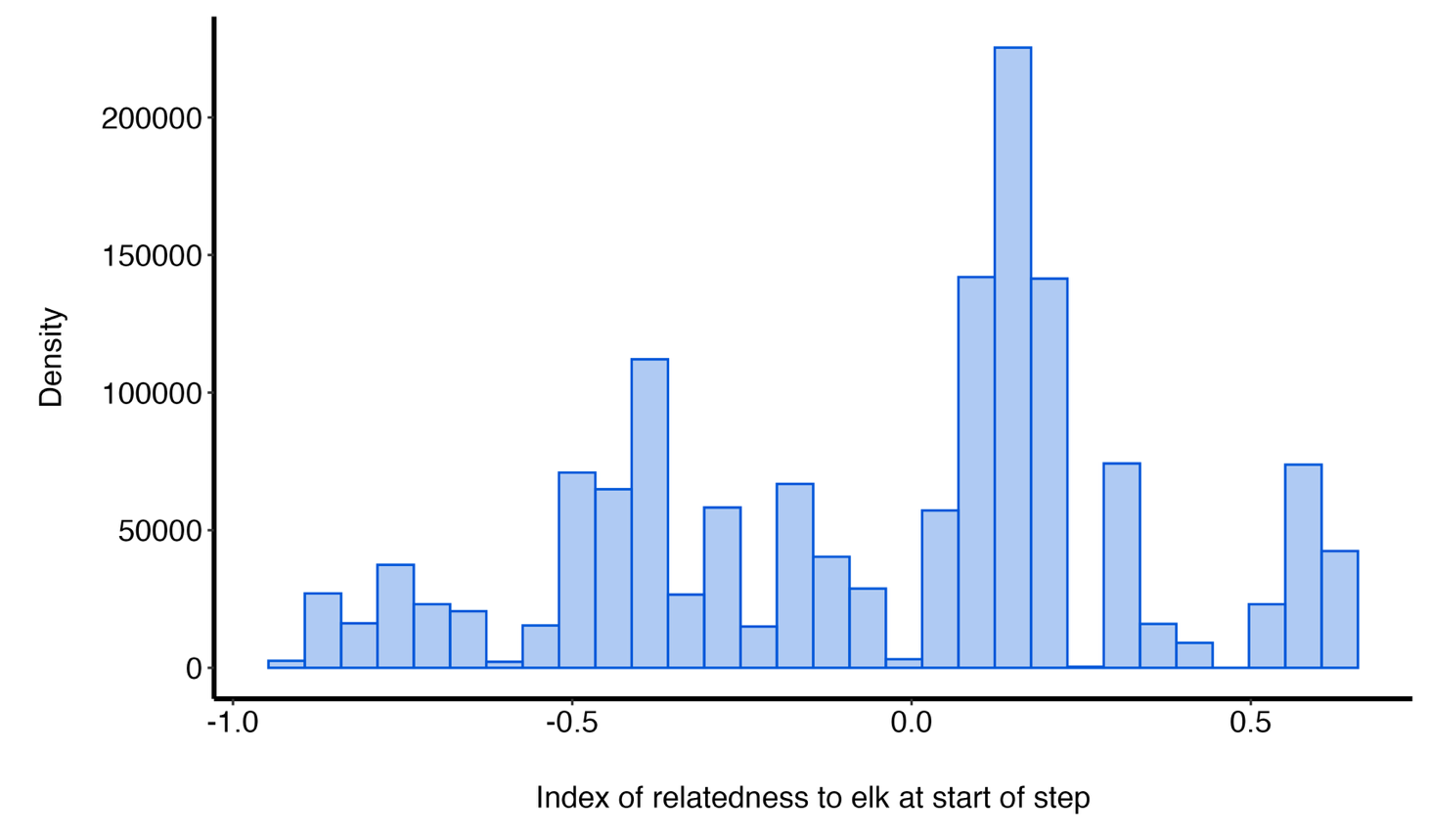

**Figure S1** Distribution of relative genetic relatedness estimates at the start of movement steps included in the inclusive fitness model.

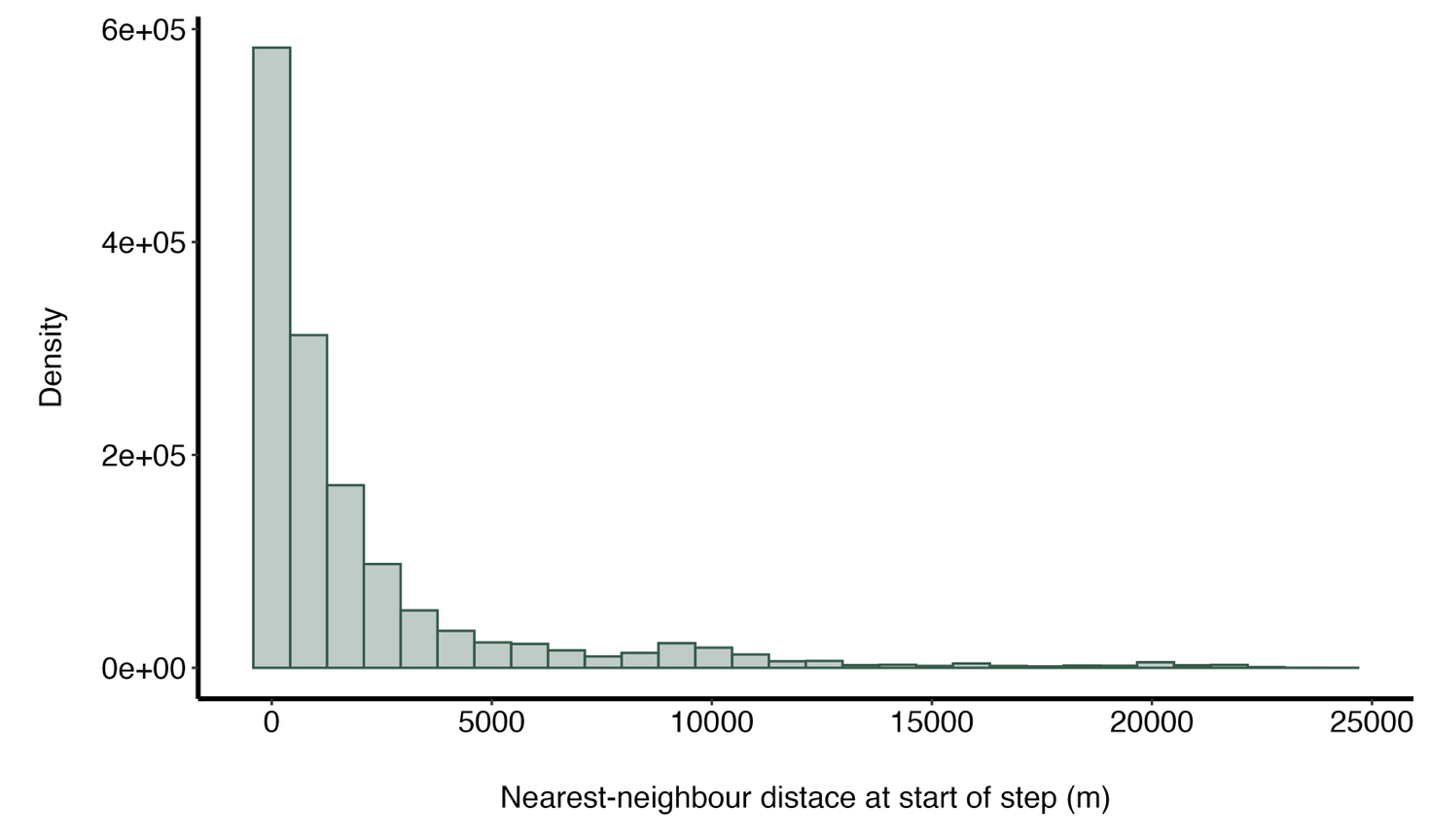

**Figure S2** Distribution of nearest-neighbour distances at the start of movement steps included in the dilution model.

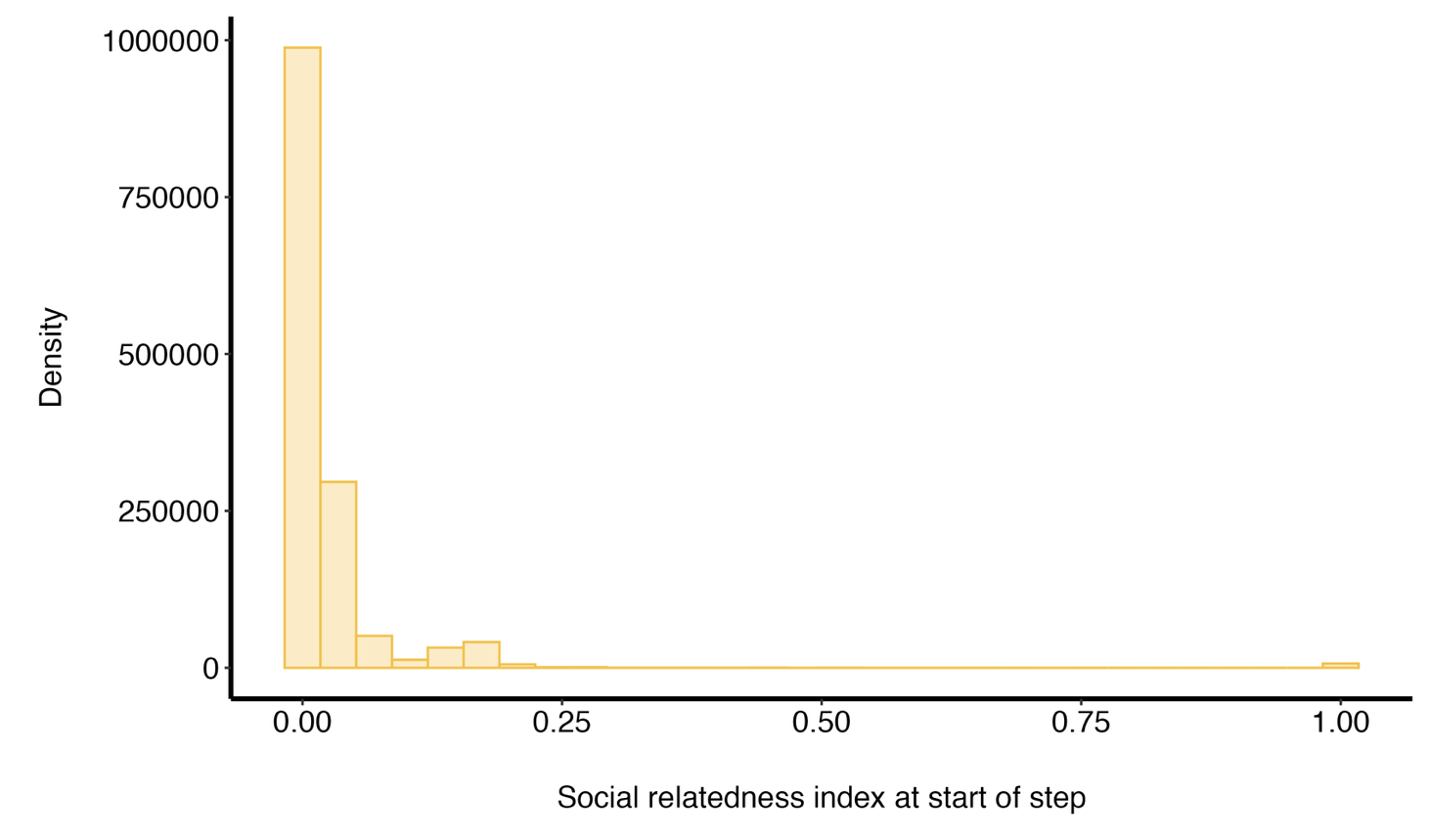

**Figure S3** Distribution of simple ratio indices at the start of movement steps included in the familiarity model.

**Table S1** Full list of model covariates included in each model

| Model |  | Fixed effect covariates | Random effects |
| --- | --- | --- | --- |
| **Familiarity** |  | Habitat | Step ID (intercept) |
|  |  | SRI | Elk ID (intercept) |
|  |  | log(SRI) x habitat | log(SRI) x habitat (slope) |
|  |  | cosine(turn angle) |  |
|  |  | log(step length) |  |
| **Dilution** |  | Habitat | Step ID (intercept) |
|  |  | NN-distance | Elk ID (intercept) |
|  |  | log(NN-distance) x habitat | log(NN-distance) x habitat (slope) |
|  |  | cosine(turn angle) |  |
|  |  | log(step length) |  |
| **Inclusive fitness** |  | Habitat | Step ID (intercept) |
|  |  | Relatedness index | Elk ID (intercept) |
|  |  | Relatedness index x habitat | Relatedness index x habitat (slope) |
|  |  | cosine(turn angle) |  |
|  |  | log(step length) |  |
| **Ecological null** |  | Habitat | Step ID (intercept) |
|  |  | cosine(turn angle) | Elk ID (intercept) |
|  |  | log(step length) |  |
